# The recovery of 12,789 genomes revealed the diversity, function, and microbial interactions of the geothermal spring microbiome

**DOI:** 10.64898/2026.03.10.710734

**Authors:** Yu-Xian Li, Yang-Zhi Rao, Ze-Wei Li, Shao-Peng Li, Yan-Ni Qu, Jia-Liang Kuang, Yong-Jian Chen, Yan-Ling Qi, Qi-Jun Xie, Wen-Sheng Shu, Meng-Meng Li, Lan Liu, Jian-Yu Jiao, Zheng-Shuang Hua, Wen-Jun Li

## Abstract

Geothermal springs, characterized by extreme physicochemical conditions, represent ecologically and evolutionarily significant habitats that foster unique microbial communities and drive adaptive evolutionary processes. Despite their importance, the complex microbial interactions and underlying mechanisms governing community assembly in these environments remain poorly understood. In this study, we conducted systematic sampling across 49 geothermal springs in Tengchong, Yunnan, over a six-year period (2016-2021), and performed metagenomic sequencing on 152 samples. We successfully reconstructed 12,789 non-redundant microbial genomes, revealing an exceptionally high level of phylogenetic and functional diversity within the spring microbiomes. Our analyses demonstrate that pH and temperature are the primary deterministic drivers shaping both microbial species composition and functional potential, thereby segregating the communities into three distinct groups: acidic, hyperthermal, and thermal. Furthermore, ecological network analysis revealed that extreme environmental conditions significantly alter network topology, resulting in less complex but more efficient microbial interaction networks. Collectively, this study provides a comprehensive resource and mechanistic insights into the microbial diversity, community structure, and species interactions in geothermal spring ecosystems.

## Introduction

Microorganisms are important participants in the biogeochemical process across all ecosystems on Earth^1–4^. Consequently, investigating microbial species and functional diversity is vital to understanding both microbial community dynamics and the biogeochemical cycle within ecosystems. Geothermal springs, as typical high-temperature terrestrial environments with distinct geochemical characteristics, form confined “islands” that harbor specialized and potentially ancient microbes^5–7^. Previous cultivation studies and molecular surveys have revealed geothermal springs featured bacteria such as Aquificae, Proteobacteria, Cyanobacteria, Chloroflexi, and Thermus^8,9^; as well as archaea such as Crenarchaeota, Korarchaeota, and Nanoarchaeota^8–11^. Due to their geological similarity to early Earth and relatively simple microbial community structures, geothermal springs serve as ideal model systems for understanding microbial community structure, function, interactions, and assembly processes in ancient environments^8,10,12–14^. In particular, geothermal springs in Tengchong, Yunnan, China, which are magmatic-hydrothermal systems with a legacy of Cenozoic volcanic activities^15^, nurture a notable diversity of acidic geothermal springs and unique microbial diversity compared to other high-temperature systems^15–17^. Despite these advances, the microbial interactions in geothermal springs remained largely unexplored.

Disentangling ecological drivers that govern microbial community establishment and maintain biodiversity is both essential and demanding. It is well documented that both biotic and abiotic factors influence the distribution of microbes, as they interact not only with other sympatric microbes but also with the environments they reside in^18,19^. Constructing co-occurrence networks is an effective approach for biologically investigating relationships and interactions among microorganisms within complex environments^20,21^. Interpreting the network structure through various parameters and identifying key species and functional groups can provide insights into how these microbes stabilize and support community establishment^22,23^. Previously, network inference largely relied on single-marker gene-based amplicon sequencing. While this approach is simple and cost-effective, it has notable limitations, such as limited taxonomic resolution, amplification biases, and a lack of functional information. The advent of massively parallel DNA sequencing has revitalized microbial ecology, enabling more in-depth studies of microbial interactions and ecological processes that govern the community assembly. This advancement allows for a more comprehensive investigation of microbial communities at both the species and functional levels.

Based on the 12,789 MAGs reconstructed from in-depth genome-resolved metagenomic sequencing, here we uncovered unexpectedly high microbial diversity in geothermal springs, spanning 110 archaeal and bacterial phyla, with approximately 93% of identified genera being newly discovered. Strong environmental heterogeneity caused the segregation of samples into three distinct groups, where the dominant microbes in each group were also the key functional components. By considering the environmental physicochemical characteristics and the distribution patterns of microbial communities, we performed an in-depth analysis integrating molecular ecological networks (MENs). This integrative framework disclosed how biotic and abiotic factors sustain the biodiversity in geothermal spring ecosystems. Our study offers comprehensive insights into the diversity, function, and microbial interactions in hot spring ecosystems, bridging gaps in understanding these complex microbial species-functional dynamics.

## Results and discussion

### Diversity and novelty of the hot spring microbiome

A total of 152 samples covering 49 hot springs located in Tengchong (Yunnan, China) were collected and metagenomically sequenced, spanning six years (2016-2021) (Fig. S1a). The sampling sites displayed tremendous environmental heterogeneity across space and time in physiochemical parameters (Fig. S1b, Supplementary Table 1), especially covering a wide range of temperatures (23-99°C) and pH values (2.0-9.73) (Fig. S1c). With manual curation, we successfully reconstructed 20,849 MAGs from all 152 metagenomes (Fig. 1a). Among them, 5,990 (28.7%) high-quality (completeness ≥ 90%, contamination < 5%) and 6,799 medium-quality (completeness ≥ 50%, contamination < 10%) were kept for further analyses. Genomic information statistics showed the high reliability of the final set of 12,789 MAGs (Fig. S2). Remarkably, our tremendously diverse genome set was unexpectedly comparable to the genome set recovered from over 20-fold of soil metagenomes (3,304) in a recent study^24^. Regarding genomes with completeness >90%, and contamination <5%, the number of genomes in our study (5,990) even slightly surpassed (5,184). Taxonomic assignment based on the Genome Taxonomy Database (GTDB r207)^25^ identified 2,944 archaeal and 9,845 bacterial genomes (Supplementary Table 2), clustering them into 3,176 non-redundant species at 95% average nucleotide identity (ANI)^26,27^. The phylogenetic tree of these 3,176 RMAGs revealed unexpected diversity and novelty among microbes inhabiting the geothermal spring ecosystem (Fig. 1b)^28^. All RMAGs spanned 12 archaeal and 98 bacterial phyla, covering 63.2% of all known microbial phyla (GTDB r207). Among them, five phyla, including Thermoproteota, Patescibacteria, Bacteroidota, Chloroflexota, and Proteobacteria, each contained more than 200 RMAGs. Surprisingly, the phylogenetic diversity of geothermal spring samples also exceeded the phylogenetic diversity of soil at the phylum level (11 archaeal phyla and 88 bacterial phyla) in a previous study^24^. This could be a result of the immense environmental heterogeneity in our geothermal spring sampling sites (Fig. S1b and c), and the “nested” microbial diversity of non-extreme environments from the long evolutionary history of the terrestrial geothermal spring microbiome^13^. Moreover, RMAGs in this study exhibit remarkable taxonomic novelty, identifying two new phyla, 11 new classes, 62 new orders, 213 new families, 721 new genera, and 1,558 new species which account for 85.1% of the total diversity of the species level (Fig. 1c). Overall, this dataset represents a tremendous and comprehensive prokaryotic genomic resource from geothermal spring ecosystems, revealing a large amount of uncharted “microbial dark matter”.

**Fig. 1.**
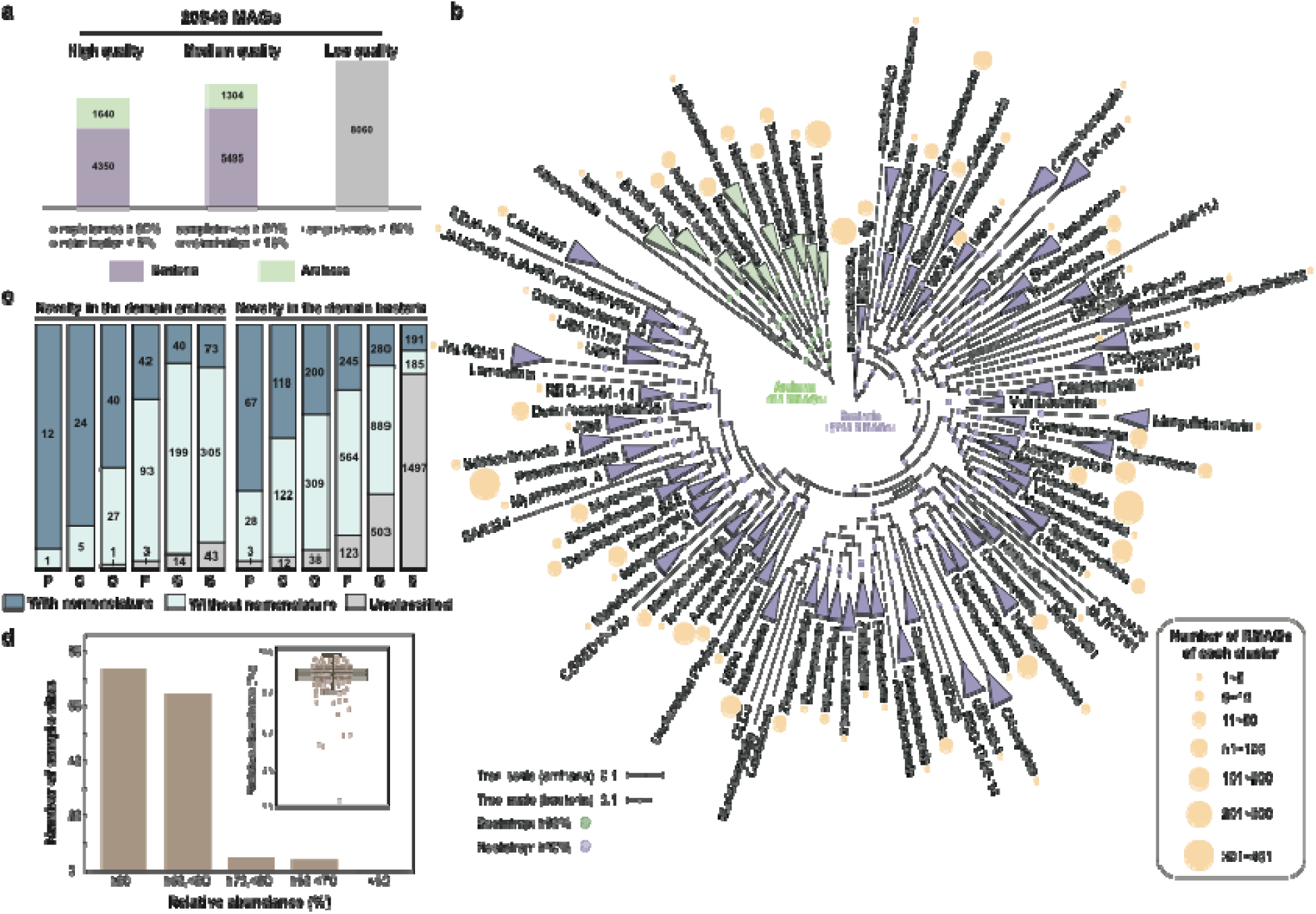
Metagenomic genome reconstruction of geothermal springs in Yunnan. **a**, Genomic quality of all 20,849 genomes reconstructed from 152 geothermal spring metagenomes. **b**, Phylogenetic distribution of 3,176 RMAGs recovered in this study. The best-fit model of the archaeal and bacterial phylogenetic tree was LG+F+R10. Clades with a bootstrap value ≥ 90% were marked by green and purple dots for the archaeal and bacterial trees respectively. Archaeal classes and bacterial phyla without nomenclature were shown in red text. The orange circles represented the number of RMAGs of each archaeal class and bacterial phylum. **c**, Stacked bar plot for novelty quantification of all archaeal and bacterial genomes. **d**, Boxplot showed the relative abundance of 3,176 RMAGs in 152 samples. The bar plot counted the number of sample sites with different ranges of relative abundance of RMAGs.

*rpS3* gene or 16S ribosomal RNA gene was often used to represent the composition of the microbial community^26,22^. However, the short sequence length and potential misassignment of the *rpS3* sequence to the associated bin limit the classification power of *rpS3*. In this study, we implemented a genome-coverage-based approach to calculate the relative abundance of microbes (Fig. S1), directly linking community compositions with genome-resolved microbial functions and thereby avoiding less accurate predictive methodologies (e.g., PICRUST)^29^. The reads mapped to 3,176 RMAGs covered an average of 88% of the entire microbial community across 152 samples (Fig. 1d, Supplementary Table 3), indicating that reconstructed RMAGs in this study are highly representative of the *in situ* microbial communities. Apparently, microbial community compositions differ at the phylum level (Fig. 2a, Supplementary Table 4). Archaea were significantly predominant (Mann-Whitney *U* test, P < 2.2e-16, Supplementary Table 5) in geothermal springs with ultra-low pH (< 5) or ultra-high temperature (> 75 °C), including DeReTiYan (DRTY), GuMingQuan (GMQ), ShuiReBaoZha (SRBZ), ZiMeiQuan (ZMQ), and ZhenZhuQuan (ZZQ) (Fig. 2a). When pH < 5, samples such as DRTY-3, DRTY-6, and DRTY-9, were dominated by Thermoplasmatota and Thermoproteota (Fig. 2b). In hyperthermal geothermal springs (GMQ, SRBZ, ZMQ, and ZZQ), Thermoproteota and Aquificota were the most dominant archaea and bacteria, respectively (Fig. 2b and c). As for the rest of the geothermal springs with less extreme environmental conditions, communities were mainly composed of bacteria from a wide variety of phyla, with Chloroflexota, Proteobacteria, Acidobacteriota, Bipolaricaulota, and Armatimonadota being the most abundant (Fig. 2c). Despite discernible differences in community compositions among geothermal springs, metabolic profiles based on KEGG Orthology (KO) categories remain remarkably consistent across all 152 samples (Fig. 2d, Supplementary Table 6 and 7). Such patterns were coherent with previous notions that higher-level functional turnover was relatively stable compared to changes in species compositions^30,31^. Principal Coordinates Analysis (PCoA) using dissimilarity matrices of RMAG and KO abundance revealed that all samples were segregated into three distinct groups based on the pH and temperature (Figs. 2e-h, Supplementary Table 1). In comparison, the microbial communities varied little over temporal and seasonal scales (winter or summer) (Fig. S3). Therefore, the community structure of geothermal springs showed greater variations over physiochemical changes than temporally, with pH and temperature being the most vital environmental drivers. Additionally, although different samples remain consistent at higher-level functional turnover (Fig. 2d), the lower-level functional genes were more sensitive to environmental changes (Fig. 2h).

**Fig. 2.**
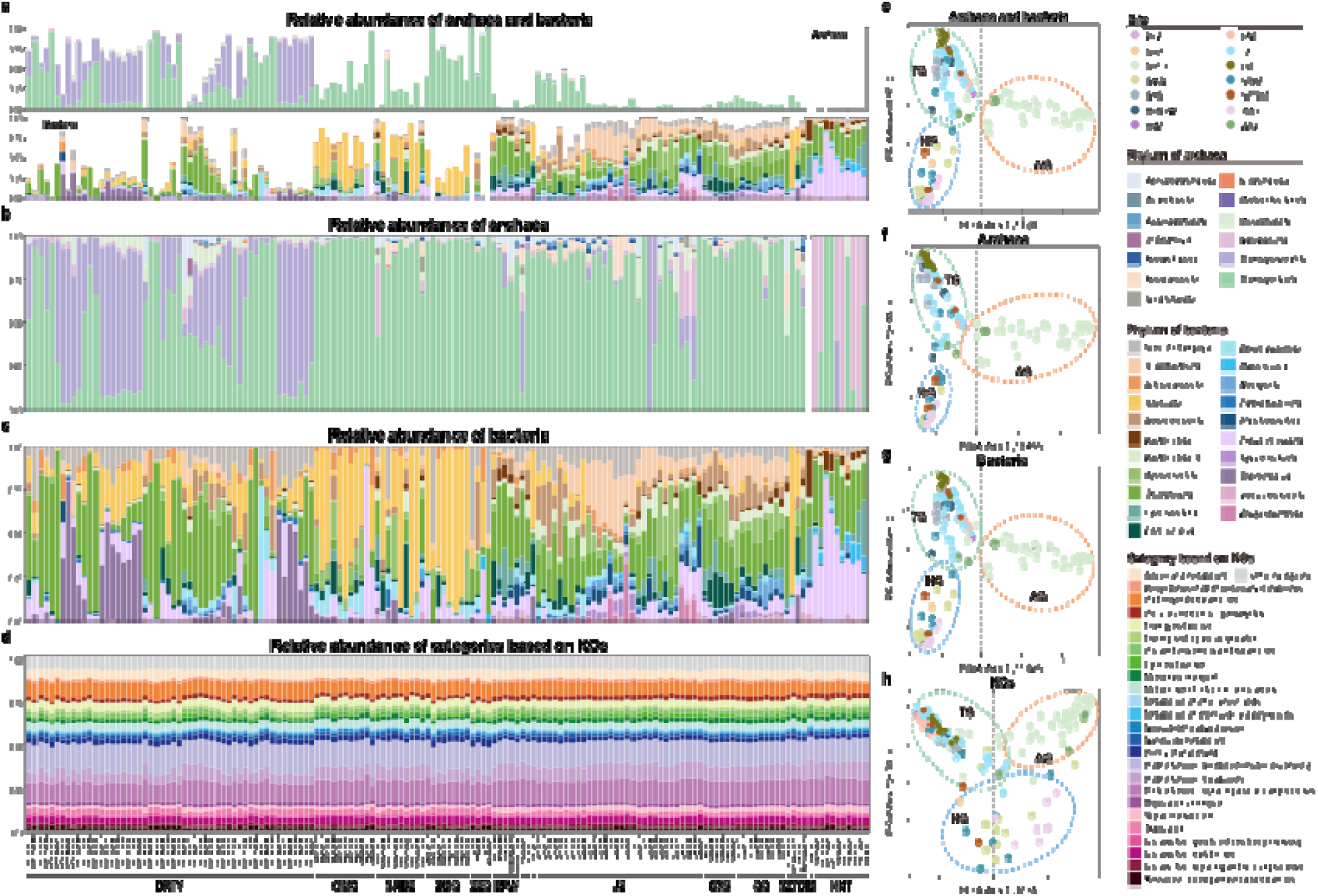
Community composition and functional profiles of 152 samples. **a,** The relative abundance of archaea and bacteria in 152 samples. **b**, The proportion of all archaea in all samples. Bars in different colors represented different classes of archaea. **c**, The proportion of all bacteria in all samples. Bars in different colors represented different phyla of bacteria. **d**, the relative abundance of KEGG standard categories (level B) in all samples. **e-f**, Principal coordinate analysis (PCoA) of all RMAGs, archaea, bacteria, and KOs in different sites based on Bray–Curtis distances.

### pH and temperature drove the species and functional community structure

To further unveil the major environmental determinants shaping community structure, we examined the relationship between environmental variables and microbial diversity and metabolic function using the Mantel test and redundancy analysis (RDA) (Fig. 3a, Fig. S4, Supplementary Table 8). Results showed that pH, Na^+^, Ca^2+^, temperature, and K^+^ were the five most prominent factors significantly correlated with community composition, with pH being the most deterministic. Combined with the PCoA results, we, therefore, divided the 152 samples into three groups based on their pH and temperature: 1) acidic group (AG): geothermal springs with pH ≤ 5; 2) hyperthermal group (HG): geothermal springs with pH > 5 and temperatures > 75°C; 3) thermal group (TG): geothermal springs with pH > 5 and temperatures ≤ 75°C (Supplementary Table 1). Such grouping pattern was further corroborated by the ANalysis Of Similarities (ANOSIM) test (R=0.679, P=0.001, Fig. 3b, Fig. S5). Interestingly, principal component analysis (PCA) based on environmental variables of these sites showed a similar grouping pattern (Fig. S6). This alignment suggests a strong ecological relationship between environmental conditions and microbial life, where specific factors consistently shape microbial communities across different samples. Comparisons of the three groups showed that microbial communities with extreme physicochemical conditions (AG and HG) exhibited significantly lower species and functional diversity (Fig. S7 and S8, Supplementary Table 9). Among the 3,176 RMAGs, only 351 were found to be shared among all three groups (Fig. S9a). Most of them were classified into Thermoproteota, Chloroflexota, and Proteobacteria (Fig. S9b). TG represents a relatively benign habitat and harbors the highest number of unique taxa (1,341), most of which belong to bacterial phyla, including 197 from Proteobacteria, 151 from Patescibacteria, and 149 from Chloroflexota (Fig. S9b). Although HG contains far fewer taxa, the majority (1,515 of the 1,598, or 98%) were also present in TG, demonstrating that HG represents a subset of the taxa detected in TG. In contrast, Thermoplasmatota, Micrarchaeota, Nanoarchaeota, and Thermotogota appeared to be more abundant in samples belonging to AG, while Aquificota were favored in hyperthermal habitats, and Thermoproteota and Aenigmatarchaeota inhabited both AG and GH (Fig. 3c). In comparison, other bacterial phyla such as KSB1, Myxococcota, and Desulfobacterota_B distributed more abundantly in TG.

**Fig. 3.**
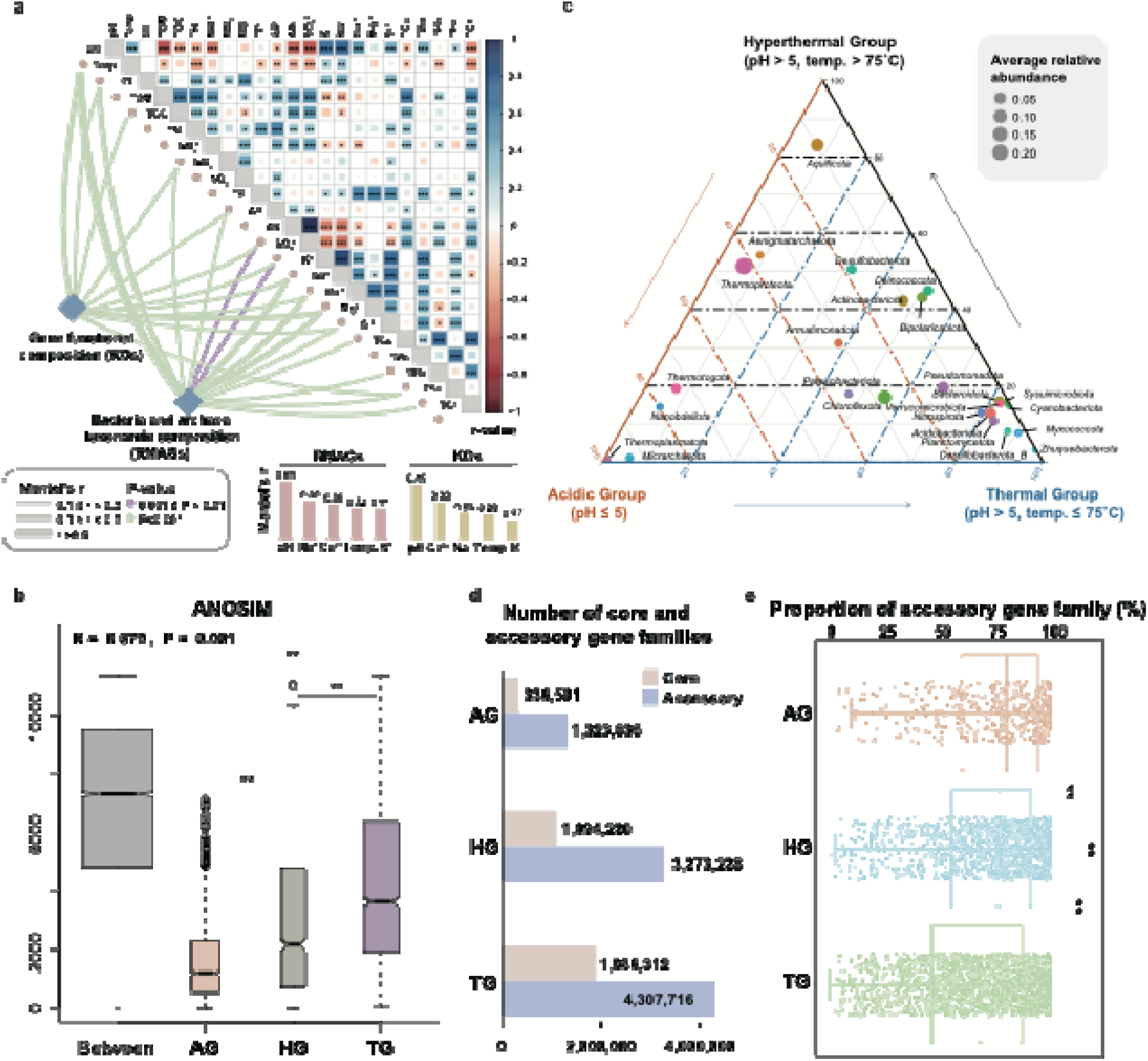
Relationship between environmental variables and species and functional composition. **a**, The heatmap showed the pairwise correlations of multiple environmental variables with the color gradient denoting Pearson’s correlation coefficients. The edges represented the correlations between community structure and environmental variables. Edge width corresponds to Mantel’s r value, and edge color corresponds to Mantel’s P value. Abbreviation: Temp., Temperature; Cl^−^, Salinaty; TOM, Total organic matter; TOC, Total organic carbon; TN, Total nitrogen; NH_4_^+^, Ammonia; NO_3_^−^, Nitrate; NO_2_^−^, Nitrite; TP, Total phosphorus; AP, Available phosphorus; AS, Available sulfur; SO_4_^2-^, Sulfate; K^+^, Exchangeable potassium; Na^+^, Exchangeable sodium; Ca^2+^, Exchangeable calcium; Mg2+, Exchangeable magnesium; Si^2+^, Effective silicon; TCu, Total bronze; TZn, Total zinc; TPb, Total lead; TFe, Total iron; TCr, Total chrome. **b**, Analysis of similarities (ANOSIM) plot showed the dissimilarity of the three groups. The width of the bar is directly proportional to the sample size. The “**” represented the significant difference (P = 0.001) between every two groups based on the pairwise ANOSIM analysis of the three groups. **c**, The ternary diagram of the relative abundance of microorganisms in three groups. The top 25 microorganisms with high average relative abundance in all 152 samples were shown in the ternary diagram. **d**, The number of core and accessory gene families of each group. **e**, Boxplot showed the proportion of accessory gene families of each group. The “**” represented the P value < 0.001 in the Wilcoxon rank sum test.

To better compare the intraspecific functional diversity of three geothermal spring groups, functional annotation was conducted on all MAGs, classifying all genes in each genome family into core and accessory genes according to their functions (see Methods, Supplementary Table 10). Among the three groups, TG had the highest number of core and accessory gene families, followed by HG and AG (Fig. 3d). This pattern likely reflects the species diversity of the three groups (Fig. S7), as higher microbial diversity typically corresponds to a larger gene pool. However, the proportion of accessory gene families was significantly higher in AG, followed by HG and TG (Fig. 3e). It is well documented that accessory genes are more dynamic, frequently entering and leaving the genome through mechanisms such as horizontal gene transfer, which enhances genetic diversity and adaptability^32^. This dynamism is likely associated with the need to acquire genes for adaptation to acidic or hyperthermal environments. By looking into the detailed functions of accessory genes, AG and HG possessed significantly more genes involved in genetic information processing (Fig. S10). Notably, approximately 73.2% (AG) and 74.5% (HG) of accessory gene families were unannotated or unclassified (Supplementary Table 11). We conjecture that species in AG and HG may rely on these unknown accessory genes to adapt to their extreme environments, necessitating further investigation into their functions.

### Species and functional network patterns differed in three environmental groups

Based on the relative abundances of 3,176 RMAGs in all samples, Molecular Ecological Networks (MENs) were built separately for the three groups to reveal the interactions among microorganisms^22,33,34^ (Fig. 4a, b, and c). Generally, all three networks had “non-random”, “scale-free” and “small-world” properties and modular structures (Supplementary Table 12)^22^. The network size (total number of nodes) and network connectivity (total number of links)^35,22^ in the three MENs demonstrated a decreasing trend as environmental conditions became harsher, with TG (343 nodes linked by 1,243 edges) having the largest MEN but AG (84 nodes linked by 321 edges) and HG (172 nodes linked by 553 edges) possessing much smaller MENs. Despite having a smaller MEN, AG exhibited a higher average degree (or average connectivity, i.e., average links per node), average clustering coefficient (the extent to which nodes are clustered), and connectedness (the proportion of realized links in all possible ones), indicating a higher level of network clustering within the community (Fig. 4d, e, and g, Supplementary Table 13). Therefore, the neutral process might be less prevalent when pH decreases^35^. Compared to TG, the higher magnitude of local clustering and shorter average path distance in AG and HG suggested a faster response to environmental disturbances in geothermal springs with physiochemical extremes (Fig. 4e and f), which are also hallmarks of small-world networks. All MENs in the three groups were highly modular, with 19 large modules (modules with ≥ 5 nodes) accounting for 88-97.6% of the nodes. This demonstrated the higher stability and robustness of MENs in terrestrial geothermal ecosystems. It also reflected that the entire community may be organized into distinct sub-communities, with possible functional specialization occurring within each sub-community. This was especially true for the HG group, which exhibited the highest modularity (Fig. 4b). Interestingly, the physiochemical extremes (low pH and high temperature) have a pronounced effect on the composition of MENs. In AG, nodes with higher degrees were mostly assigned to archaea, including Thermoplasmatota, Micrarchaeota, and Thermoproteota. These nodes exhibited strong interactions with those from other phyla, such as Aenigmatarchaeota and Desulfobacterota (Fig. 4a, Supplementary Table 10). In HG, high-degree nodes were mainly classified into both bacteria and archaea, including Thermoproteota, Aquificota, and Deinococcota, with each module consistently dominated by microbes from the same phylum (Fig. 4b). This also corroborated the presence of possible functional specialization within different modules. The MEN of TG showed higher microbial diversity and complexity, with Bacteria playing more significant roles in the relatively benign environments (Fig. 4c).

**Fig. 4.**
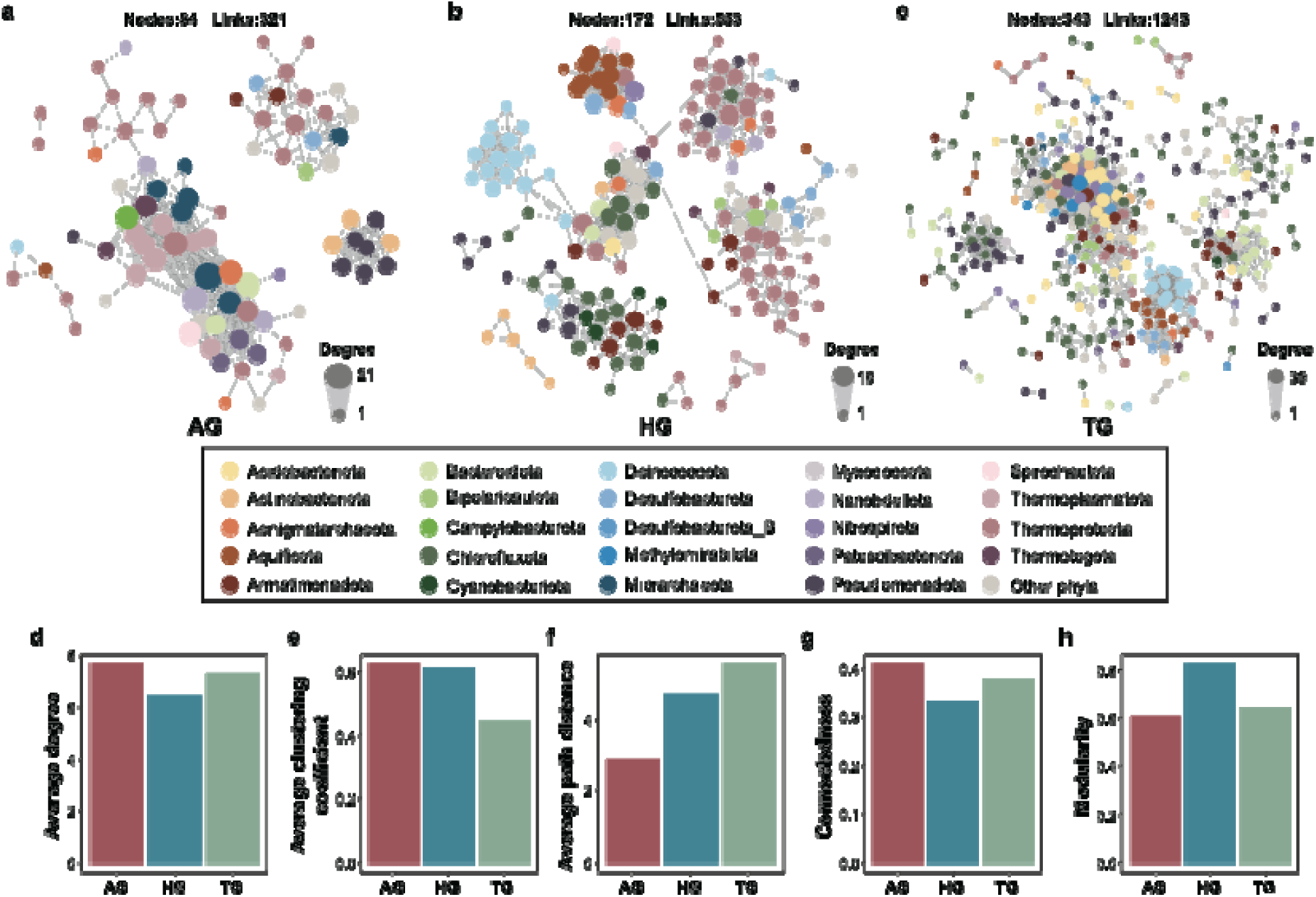
Co-occurrence microbial networks of three geothermal spring groups. **a-c**, Visualization of constructed MENs of **a**, AG, **b**, HG, and **c**, TG. The nodes were colored according to the phylum taxonomy of RMAGs. The circle size represented the nodes’ degrees, and the values listed in Supplementary Data 10. A connection represents a strong (Spearman’s correlation coefficient ≥ 0.78) and significant (P-value < 0.01) correlation. **d-h**, Important network properties of three groups, including **d**, Average degree, **e**, Average clustering coefficient, **f**, Average path distance, **g**, Connectedness, **h**, Modularity. Details of all network properties are listed in Supplementary Data 9.

To unravel the functional specialization performed by microbes in each module and potential functional interactions among different modules, as well as to establish links between microbes at both the species and function levels, we further constructed functional networks for the three groups based on the relative abundance of core functional genes (KOs) (Fig. S11-13, Supplementary Table 14). The top 50 functional genes with the highest node degree were selected to represent the core metabolic features of certain communities (Supplementary Table 15 and 16). Overall, the key metabolic processes of the three groups appeared to be markedly distinct. We observed that dissimilarity sulfate reduction (DSR) and some carbon metabolism pathways, including tricarboxylic acid cycle (TCA), pyruvate metabolism, and fermentation, played a significant role in AG, and sulfur oxidation and carbon fixation pathways, such as reverse tricarboxylic acid cycle (rTCA), dicarboxylate/4-hydroxybutyrate or dicarboxylate/hydroxybutyrate (DC-HB) cycle, and Wood-Ljungdahl (WL) pathway, were crucial in HG. As for TG, the WL pathway, rTCA, electron transport chain (ETC), TCA, glycolysis, pyruvate metabolism, fermentation, and dissimilatory nitrate reduction (DNR) were the most important functions. Such metabolic distinction in the three groups could be attributed to functional cooperation performed by microbes from different modules.

## Conclusion

This study is the first to systematically sample geothermal spring ecosystems at both spatial and temporal scales, albeit locally, to explore how environmental parameters influence the composition and function of microbial communities. From 12,789 bacterial and archaeal MAGs reconstructed from 152 metagenomes, we uncovered immense phylogenetic diversity among microbes inhabiting terrestrial geothermal springs, with both species and functional diversity primarily shaped by pH and temperature, which were divided into three groups: acid group (AG), hyperthermal group (HG), and thermal group (TG). By combining species and functional MENs, we found that networks in samples with extreme physicochemical conditions (AG and HG) were less complex but more efficient than those in relatively benign environments (TG). Microbial communities in HG formed highly modular structures to enhance cooperation, with most modules highly independent of the others. Dominant microbes define both the composition and function of the geothermal spring ecosystem, with archaea prevailing in extreme conditions, especially in AG. Altogether, our study comprehensively and thoroughly elucidates the microbial species and functional diversity, as well as microbial interactions of geothermal spring ecosystems.

## Methods

### Study sites, sampling, DNA extraction, and sequencing

Sediment samples were collected twice a year, once in summer and once in winter, from 49 geothermal springs located in Tengchong, Yunnan, China from 2016 to 2021. A total of 152 samples were collected and physicochemical parameters were measured. Community DNA was extracted and subsequently conducted for metagenomic sequencing. The detailed procedures for sample collection, DNA extraction, and metagenomic sequencing were described in the previous study^36^. Sample information, including sample images, sampling time, geographical locations, and physicochemical parameters were provided in Fig. S1 and Table S1. The total genomic DNA was sequenced using the Illumina Hiseq 4000 instrument at Beijing Novogene Bioinformatics Technology Co., Ltd (Beijing, China). Raw sequencing data for each sample was generated with an average amount of 30 Gbp (2 × 150 bp).

### Metagenomic assembly, genome binning, and curation

Raw sequencing reads were preprocessed as described in detail in the previous study^37^. The quality reads of each sample were *de novo* assembled using SPAdes^38^ (version 3.15.2) with parameters: -k 21,33,55,77,99,127 --meta. Scaffolds with lengths < 2500 bp in each assembly were eliminated for subsequent analyses. BBMap^39^ (version 38.92, http://sourceforge.net/projects/bbmap/) was used to calculate the coverage information of each scaffold with the following parameters: k=15 minid=0.97 build=1. Multiple steps were performed to obtain high-quality genomes from assembled sequences. First, genome binning was conducted by using MetaBAT^40^ (version 2.12.1), CONCOCT^41^ (version 1.1.0), and MaxBin 2^42^ (version 2.2.7) separately. DAS Tool^43^ was subsequently applied to identify the most accurate and non-redundant bins from the combined bins generated by the three tools. Reads were recruited by mapping the respective metagenomes to the selected bins using BBMap. Reads recruited from each bin underwent further reassembly using SPAdes with parameters: -k 21,33,55,77,99,127 --careful. Finally, a total of 20,849 MAGs with high- and medium-quality was retained.

Genome quality, including completeness, contaminations, and strain heterogeneity were estimated using CheckM^44^ (version 1.1.3). For bins associated with DPANN and CPR, genome quality was also assessed by calculating the occurrence frequency of 48 and 43 single-copy genes (SCGs) respectively^26^. Bins with completeness < 50% were discarded. Genome bins with contamination > 3% were decontaminated by identifying scaffolds with multiple SCGs and discordant coverage. Transfer RNAs were identified using tRNAscan-SE^45^ (version 2.0). Ribosomal RNA genes including 5S, 16S, and 23S were detected using RNAmmer^46^ (version 1.2) and BLAST^47^ (version 2.12.0+) searches against the RDP database^48^. To further improve the quality of genomes, all 16S rRNA genes were aligned using the online tool SINA (version 1.2.12) (https://www.arb-silva.de/aligner/) with default parameters for taxonomy assignments. Based on the taxonomy assignment of all genomes, archaeal genomes containing 16S rRNA assigned to bacteria or vice versa, and genomes with over four 16S rRNA gene copies were removed. After completing all the screening steps mentioned above, a total of 12790 genomes were subjected to further analyses. Dereplication was conducted on these MAGs using dRep^49^ (version 3.2.2) at a 95% ANI threshold (species level)^26^, resulting in the generation of 3,176 representative MAGs (RMAGs).

### Taxonomic, functional annotation, and pan-genome analysis of RMAGs, and

Taxonomic information of all 12,789 MAGs was determined by GTDB-Tk^28^ (version 2.0.0; GTDB release 207) with the “classify_wf” command. Taxonomy of MAGs did not match the taxonomy of the RMAG they were readjusted according to the RMAGs within each cluster. Putative protein-coding genes (CDSs) of all 12,789 MAGs were predicted using Prodigal^50^ v2.6.3 under the “-p single” mode with standard genetic code and annotated against the KEGG database with parameters: -E 1e-5 using kofam_scan^51^ (version 1.3.0). To investigate the microbial pangenome, gene families were identified from RMAGs with more than one genome. CDS predicted by Prodigal were first subject to all-to-all blast. Only blast hits with identity ≥ 90%, aligned region ≥ 100bp, and covering ≥ 50% of subject and query genes were kept and subsequently clustered into gene families using MCL^52^ v21-257 with parameter: -I 1.4^53^. The functions of each gene family were determined according to the voting of each annotated protein sequence.

### Phylogenetic analysis

The 3,176 RMAGs representing all microbes inhabiting Tengchong geothermal springs were used to reconstruct the phylogeny. Based on the recommendation by GTDB-Tk, 120 and 53 conserved marker genes were selected for reconstructing the bacteria and archaea phylogenies, respectively. Alignment profiles of concatenated marker genes generated by GTDB-Tk were extracted, and poorly aligned regions were trimmed using TrimAl^54^ (version 1.4.rev22) with parameters: -automated1. IQ-TREE^55^ (version 1.6.12) was applied to reconstruct the maximal-likelihood phylogenies of bacteria and archaea, with the best models determined to be LG+F+R10 for both bacterial and archaeal trees.

### Relative abundance calculation

The relative abundances of each RMAG in each site were calculated. Reads from each metagenome were mapped onto all 3,176 RMAGs and assembled scaffolds with lengths ≥ 500 bp using BBMap, respectively. The relative abundance of each RMAG in the corresponding sample was calculated as the proportion of reads recruited from the RMAG to the total reads recruited from all scaffolds with lengths ≥ 500 bp. Considering the bias of genome binning and mapping, false positive reads may be mismatched in mapping and generate an extremely low relative abundance of RMAGs. By comparing to the *rpS3* data, we replaced the relative abundance below 1E-4 with 0 to remove potential false positives (Supplementary Table 4). Finally, a total of 37,523 microbes with relative abundances > 0 were identified in all 152 metagenomes, which is comparable to the identified *rpS3* genes(27,436) in all metagenomes using AMPHORA^56^. This improved method may provide a reference for future microbial community ecological analysis based on metagenomic data. The relative abundance of gene *i* (*RA_Gi_*) annotated with KOs in one sample can be calculated by:

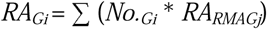

where *No._Gi_* is the number of *gene i* in *RMAG j, RA_RMAGj_* is the relative abundance of *RMAG j* in a geothermal spring sample. The relative abundance of each KO category was calculated as the sum of the relative abundances of all individual KOs within the category.

#### Network construction and analysis

All 152 samples were divided into three groups (acidic, thermal, and hyperthermal) according to their environmental parameters. MENs for each group were constructed based on community composition and function using iNAP^57^ (https://inap.denglab.org.cn). For species MENs, we used the relative abundance of 3,176 RMAGs as input. Species present in over 25% of samples within each group were kept for network construction. Then, three MENs were constructed based on Spearman correlation coefficients, and the correlation cut-off threshold was automatically determined by an RMT-based approach^22,58^. Various network properties and node properties were calculated based on the adjacent matrix using the MENA Pipeline^57^. As for functional MENs of the three groups, we selected functional genes occupied in core metabolic pathways to construct the MENs, based on the relative abundance of selected genes of RMAGs present in each species MENs. Functional genes that occupied at least 75% of samples within each group were retained for subsequent analysis. Other details regarding the construction of functional MENs were consistent with the procedures applied for species MENs. All networks were visualized using Cytoscape^59^ (v3.9.1).

#### Statistical analyses

All statistical analyses were performed in the R software (v4.2.3). Microbial community dissimilarity (beta diversity) was calculated by the Bray-Curtis index via the vegdist() function in the vegan R package (v2.6-6.1)^60^. Mann-Whitney *U* test in R package stats (v4.2.2)^61^ was used to test the difference in archaeal community between extreme geothermal springs and normal geothermal springs. Boxplot, staked bar, Venn diagram, and ternary diagram were all generated by the “ggplot2” R package (v3.4.1)^62^. The ratio differences of accessory gene families of three geochemical groups were tested by ANOVA with function aov() in R package stats (v4.2.2)^61^, and HSD post hoc all-pairwise comparisons test^63^ was conducted by function HSD.test in R package agricolae (v1.3-2)^64^. Heatmap was generated by the “pheatmap” R package (v1.0.12)^65^.

## Supporting information

Supplemeantary informations

## Code availability

All workflows for phylogeny, annotations, and comparative/evolutionary genomic analyses were deposited at Figshare (10.6084/m9.figshare.26942083). We also used the following published codes: quality_control.pl (https://github.com/hzhengsh/qualityControl), concatenate_multiple_genes.pl (https://github.com/hzhengsh/phylogeny).

## Data availability

The datasets analyzed or generated in this study can be found in our Figshare repository (10.6084/m9.figshare.26942083). All MAGs described in this study have been deposited at NCBI under the BioProject ID PRJNA544494 (the accession ID of each MAG can be found in Supplementary Table 2).

## Acknowledgments

We thank Guangdong Magigene Biotechnology Co., Ltd. China for the assistance in data analysis and the entire staff from Yunnan Tengchong Volcano and Spa Tourist Attraction Development Corporation for strong support. This work was financially supported by the National Natural Science Foundation of China (Nos. 32200002, 32400002, 91951205, and 32170014).

## Author contributions

Y.X.L., Y.Z.R., and Z.S.H. conceived the study. L.L., M.M.L., J.Y.J., Y.X.L., Y.Z.R., and Z.S.H. collected the hot spring samples. L.L., M.M.L., Y.X.L., Y.Z.R., Y.L.Q., Y.N.Q., and Q.J.X. measured physicochemical parameters and DNA extraction. Y.X.L., Y.Z.R., Z.W.L., S.P.L., J.L.K., Y.J.C., Z.S.H., Y.L.Q., and Y.N.Q. performed the metagenomic analysis, genome binning, functional annotation, and ecological analysis. Y.X.L., Y.Z.R., Z.W.L, S.P.L., Z.S.H., W.J.L., and W.S.S. wrote the manuscript. All authors discussed the results and commented on the manuscript.

