## Supplementary material for "The recovery of 12,789 genomes revealed the diversity, function, and microbial interactions of the geothermal spring microbiome": Supplemeantary informations

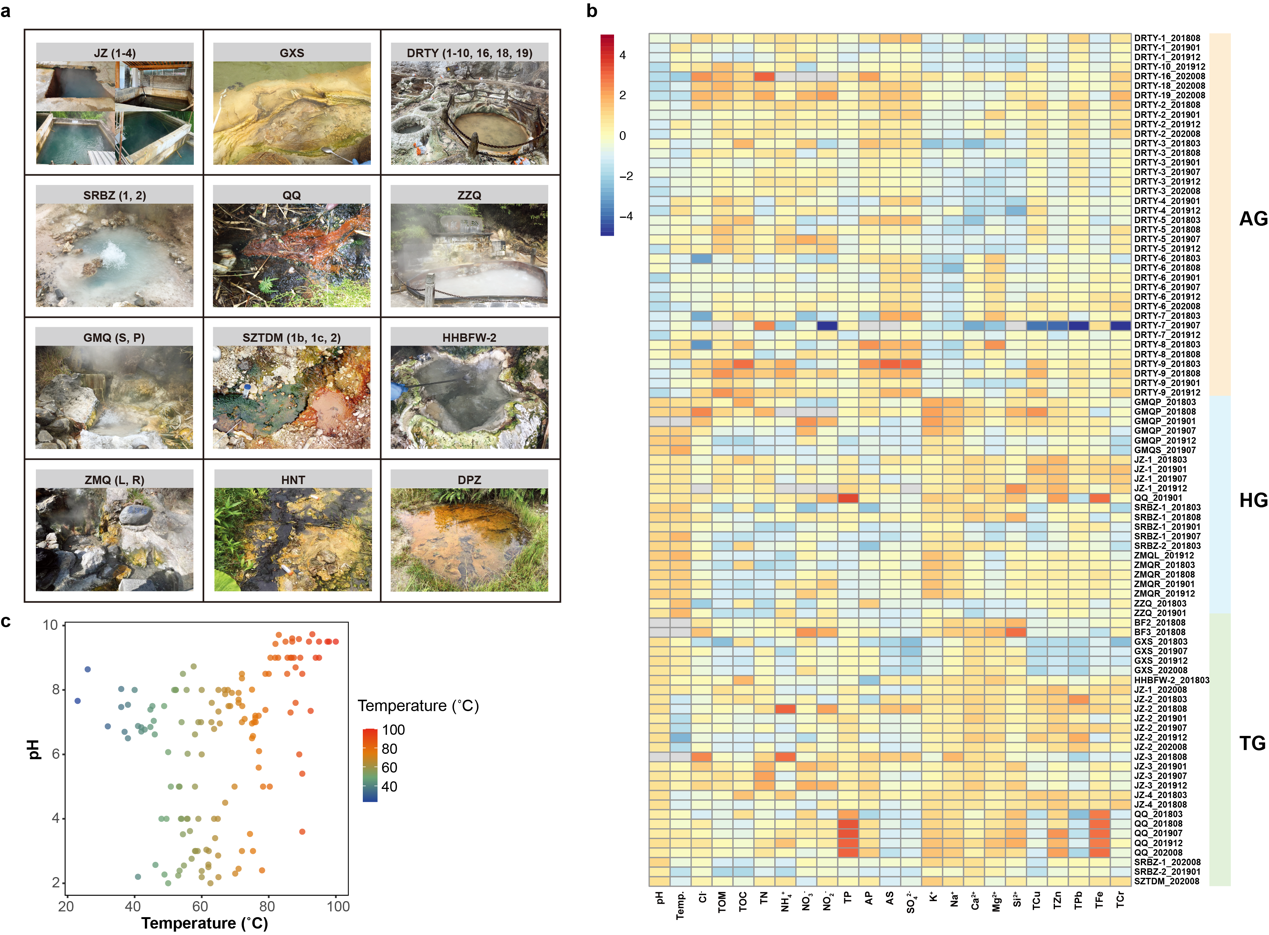

**Fig. S1. Sample sites of geothermal springs in Tengchong, Yunnan, China.** We sampled 152 metagenomes from 49 sample sites, including JZ (JinZe)-1, JZ-2, JZ-3, JZ-4, GXS (GongXiaoShe), DRC (DiReChi), DRTY (DiReTiYan)-1, DRTY-2, DRTY-3, DRTY-4, DRTY-5, DRTY-6, DRTY-7, DRTY-8, DRTY-9, DRTY-10, DRTY-16, DRTY-18, DRTY-19, SRBZ (ShuiReBaoZha), SRBZ-1, SRBZ-2, QQ (QiaoQuan), ZZQ (ZhenZhuQuan), GMQS (GuMingQuanShang), GMQP (GuMingQuanPool), SZTDM (ShiZiTouDuiMian), SZTDM-1b, SZTDM-1c, SZTDM-2, BF (BianFu)-2, BF-3, HHBFW (HeHuaBianFuWa)-2, HMZ (HaMaZui), ZMQL (ZiMeiQuanLeft), ZMQL (ZiMeiQuanRight), ZZQ (ZhenZhuQuan), ZZQ-2, DPZ (DaPingZi)-2, DPZ-5, HNT (HeiNiTan)-1, HNT-1-1, HNT-1-2, HNT-2, HNT-3, HNT-4, HNT-5, HNT-7, HNT-8, HNT-10. **(a)** Pictures of some representative sample sites. **(b)** Heatmap of physicochemical parameters of different geothermal spring samples. **(c)** Scatter plot of pH and temperature of all geothermal spring samples.

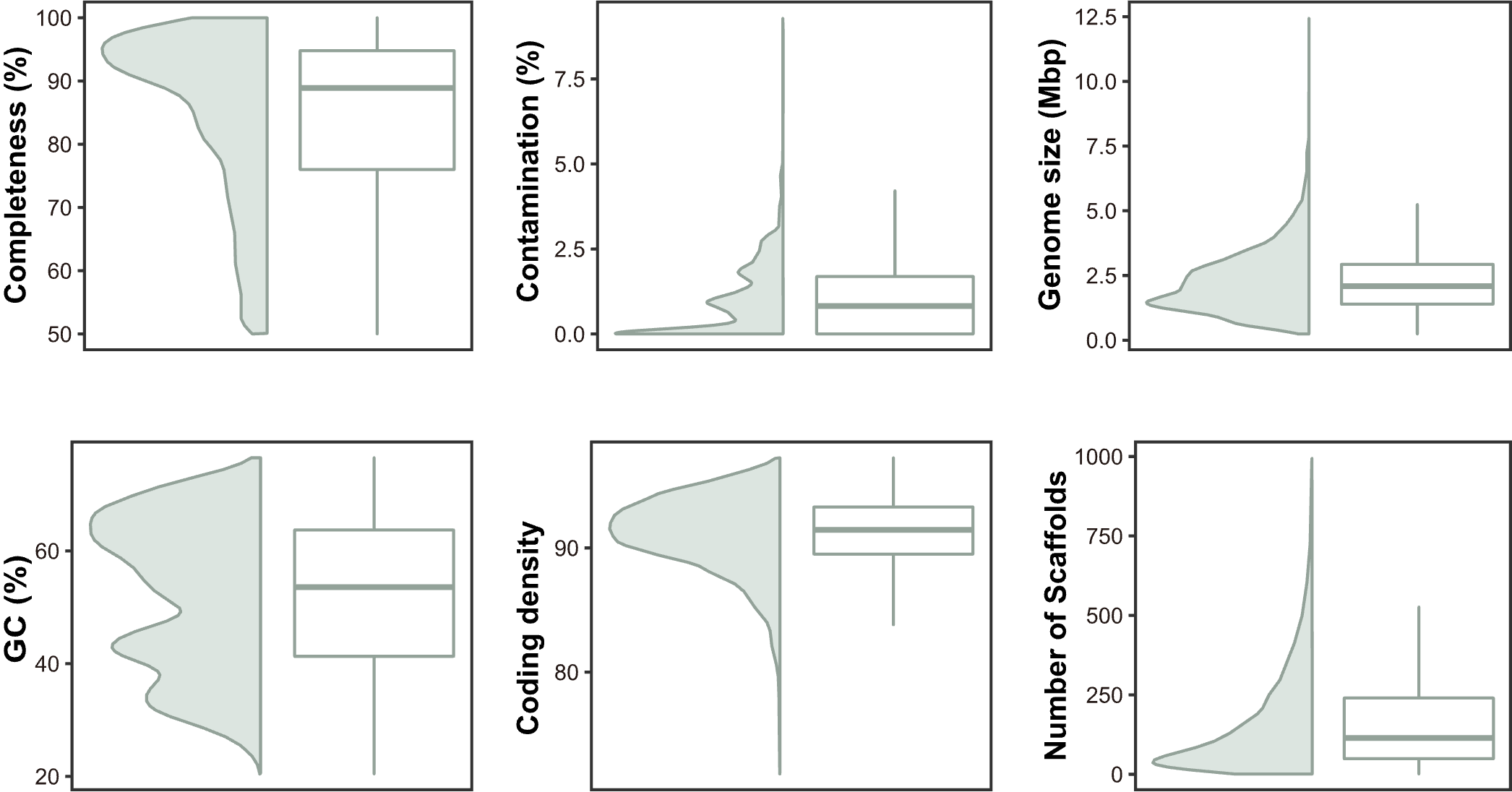

**Fig. S2. The basic genomic information of all high-quality and medium-quality genomes.** Violin plots and box plots showed completeness, GC content, contamination, coding density, genome sizes, and number of scaffolds for all 12,789 genomes.

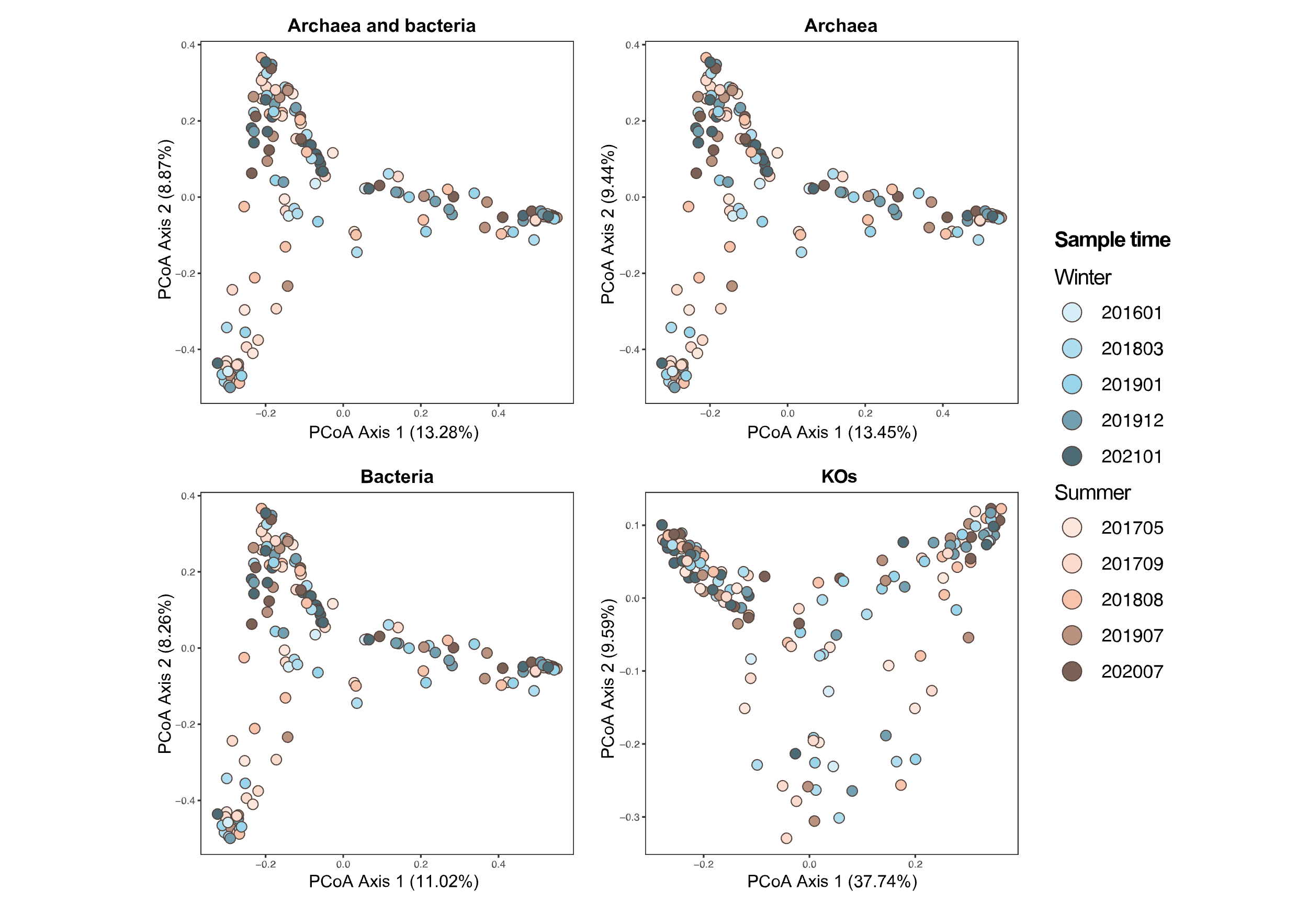

**Fig. S3. Principal Coordinates Analysis (PCoA) of all RMAGs, archaea, bacteria, and KOs in different years based on Bray–Curtis distances.**

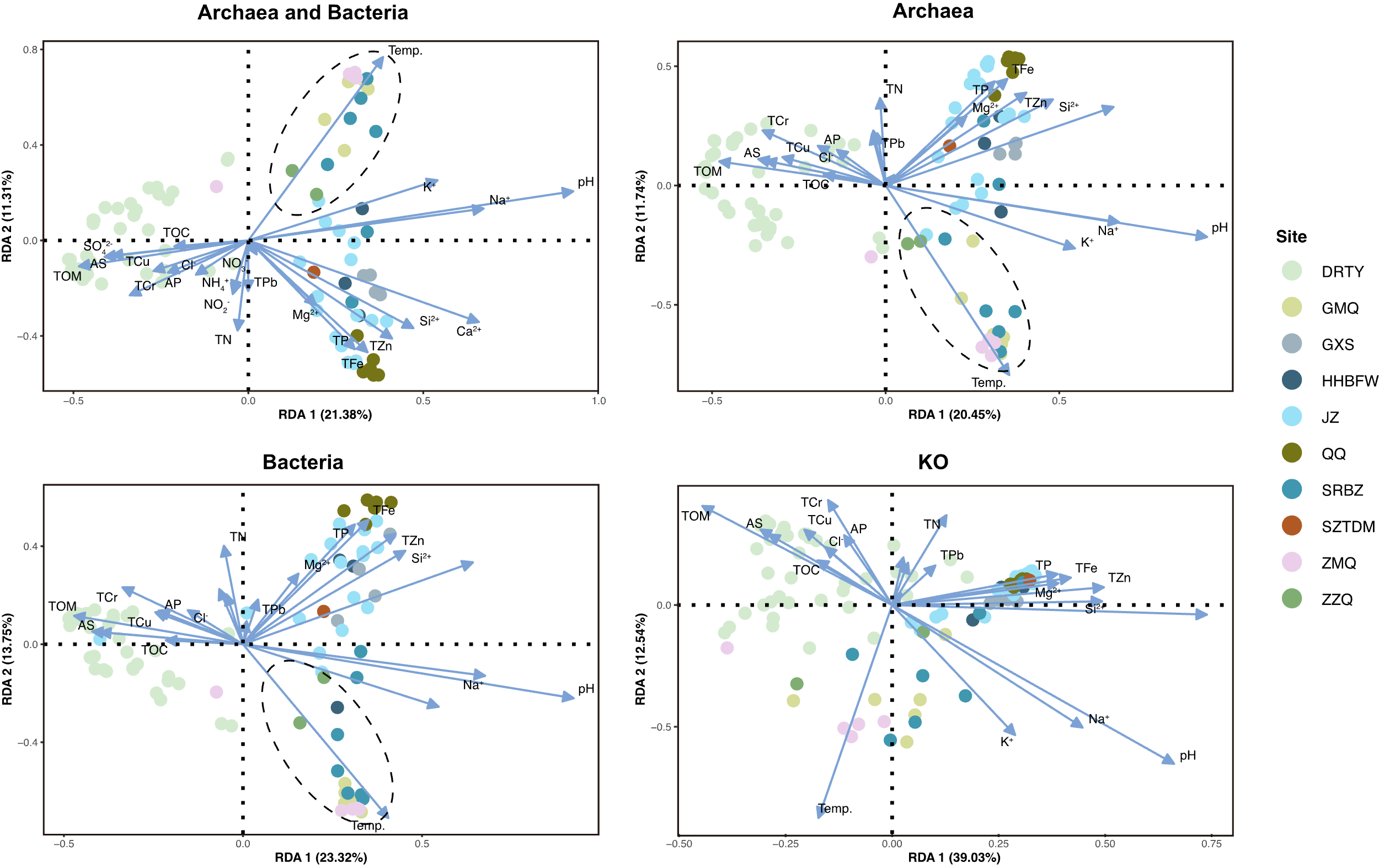

**Fig. S4. Redundancy analysis (RDA) of the relationship between the environmental variables and the microbial community composition.** Solid circles represented the geothermal spring samples. Arrows indicated the direction and magnitude of variables. 89 geothermal spring samples with consistent numbers of physicochemical parameters were selected to conduct the RDA analysis.

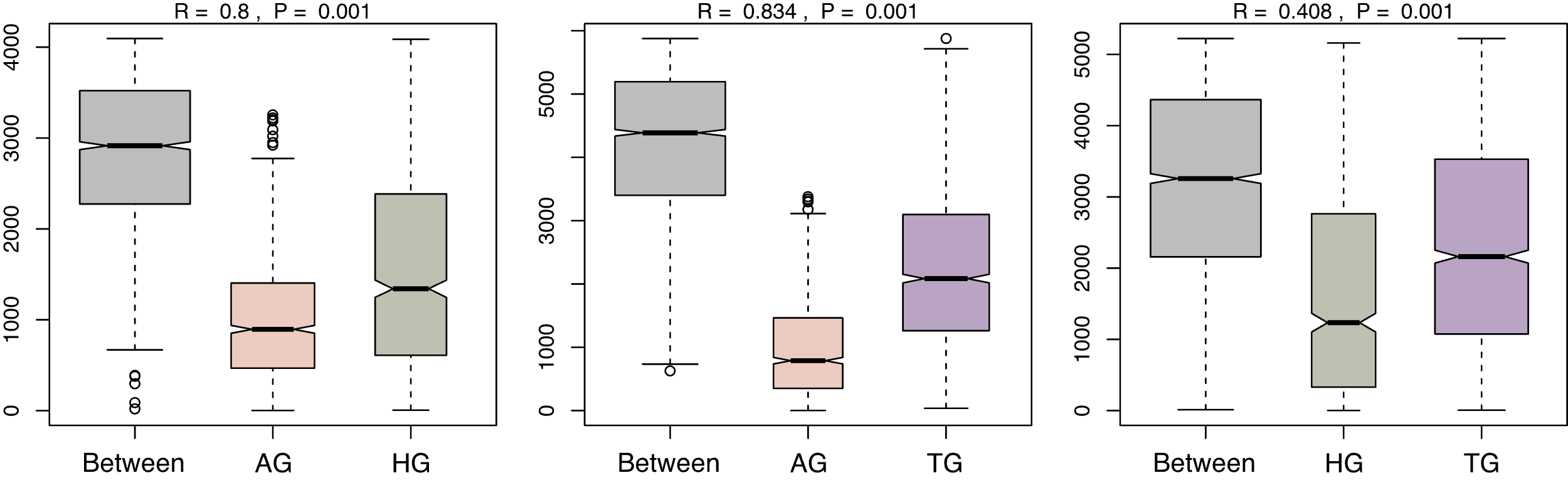

**Fig. S5.** **Analysis of similarity (ANOSIM) plot of pairwise of AG, HG, and TG.**

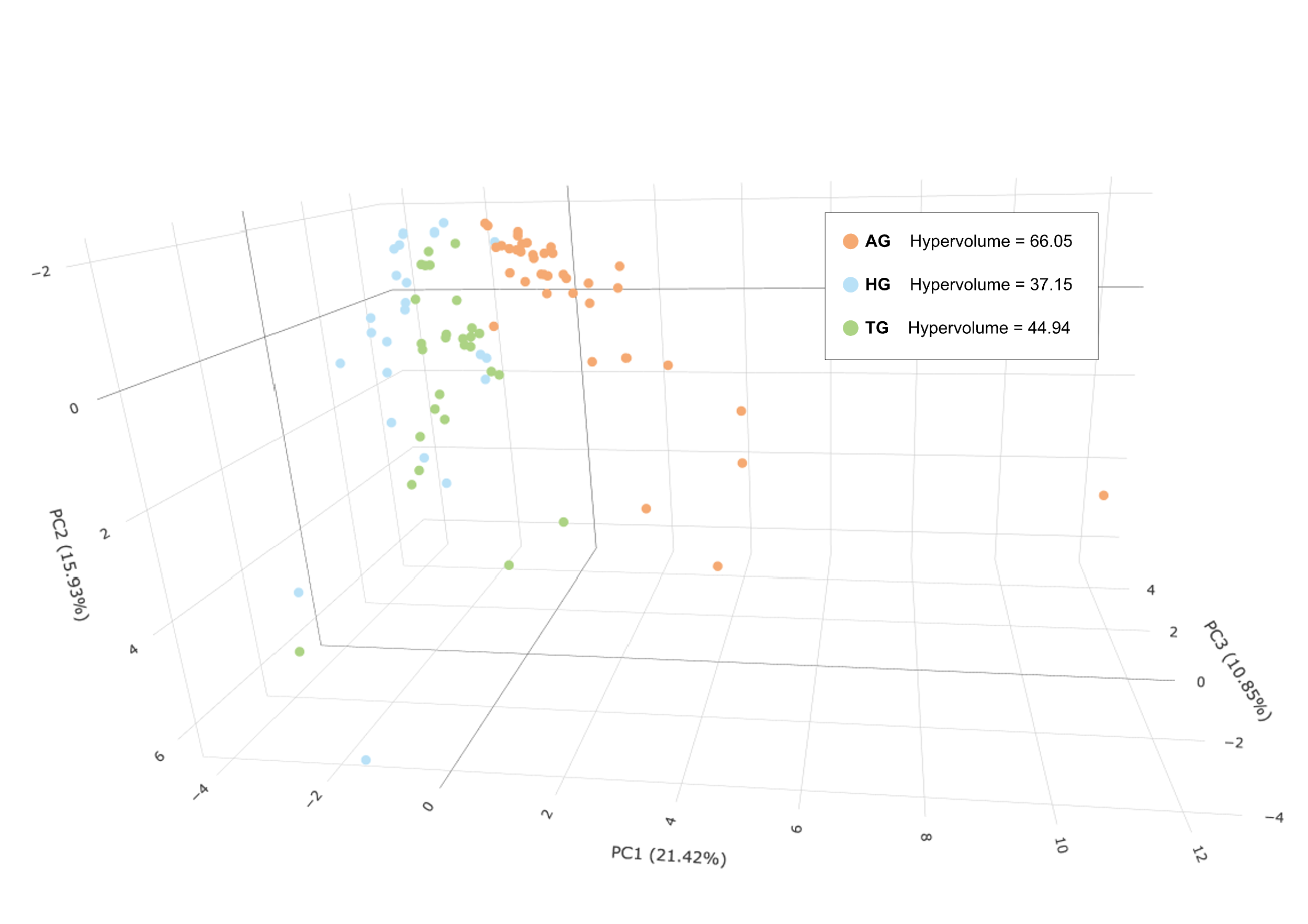

**Fig. S6. Principal component analysis (PCA) of environmental variables of Fig. S2.** The three-dimensional graph was visualized by PC 1-3 from the results. Dots with different colors, including orange, blue, and green, indicate AG, HG, and TG, respectively. Results of hypervolume computed from PC 1-3 were noted in figure legends.

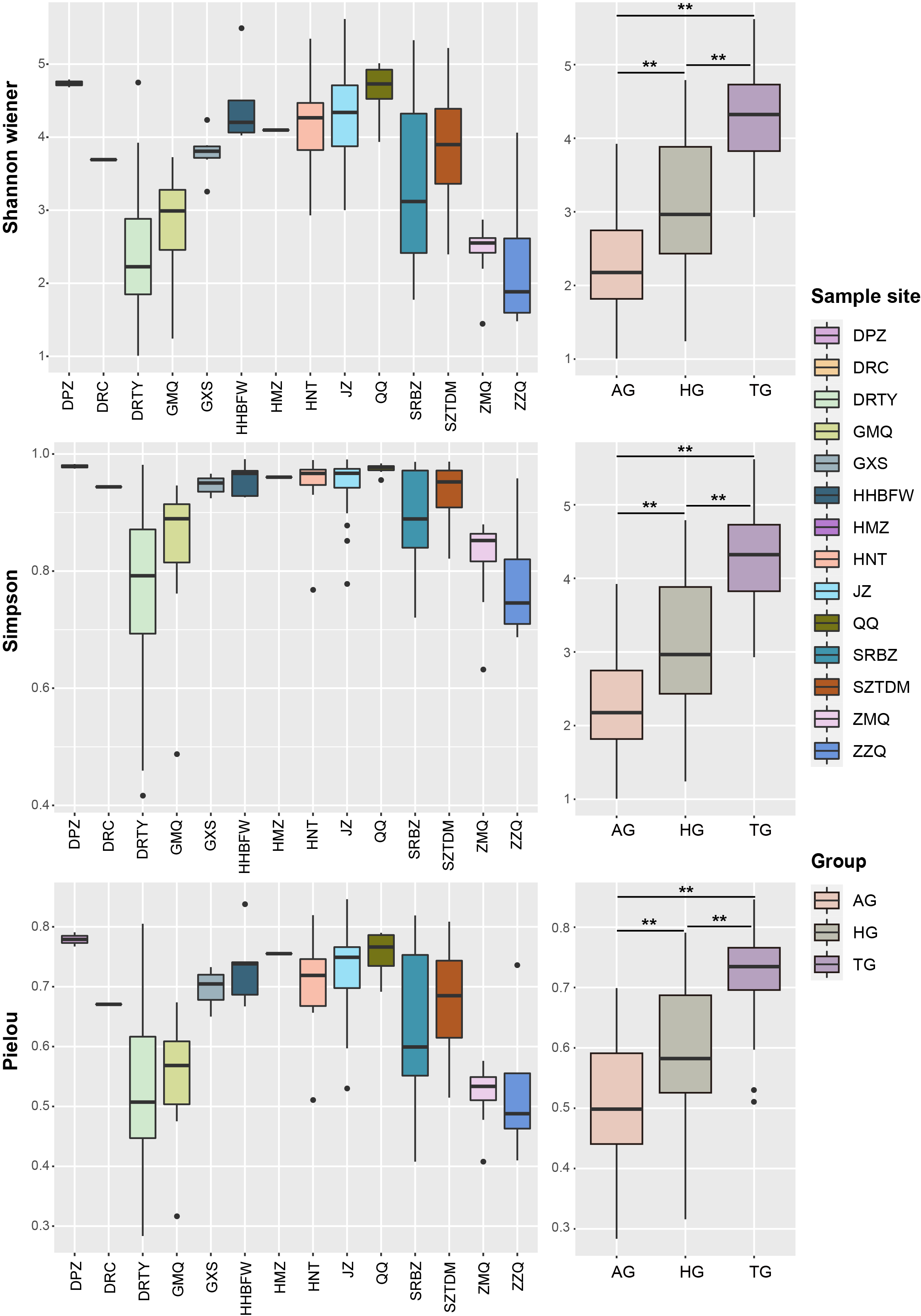

**Fig. S7. Boxplot of alpha diversity of different sites and groups.** The significant difference among the three groups was tested using the Mann-Whitney *U* test. The “**” represented the significance ≤ 0.001.

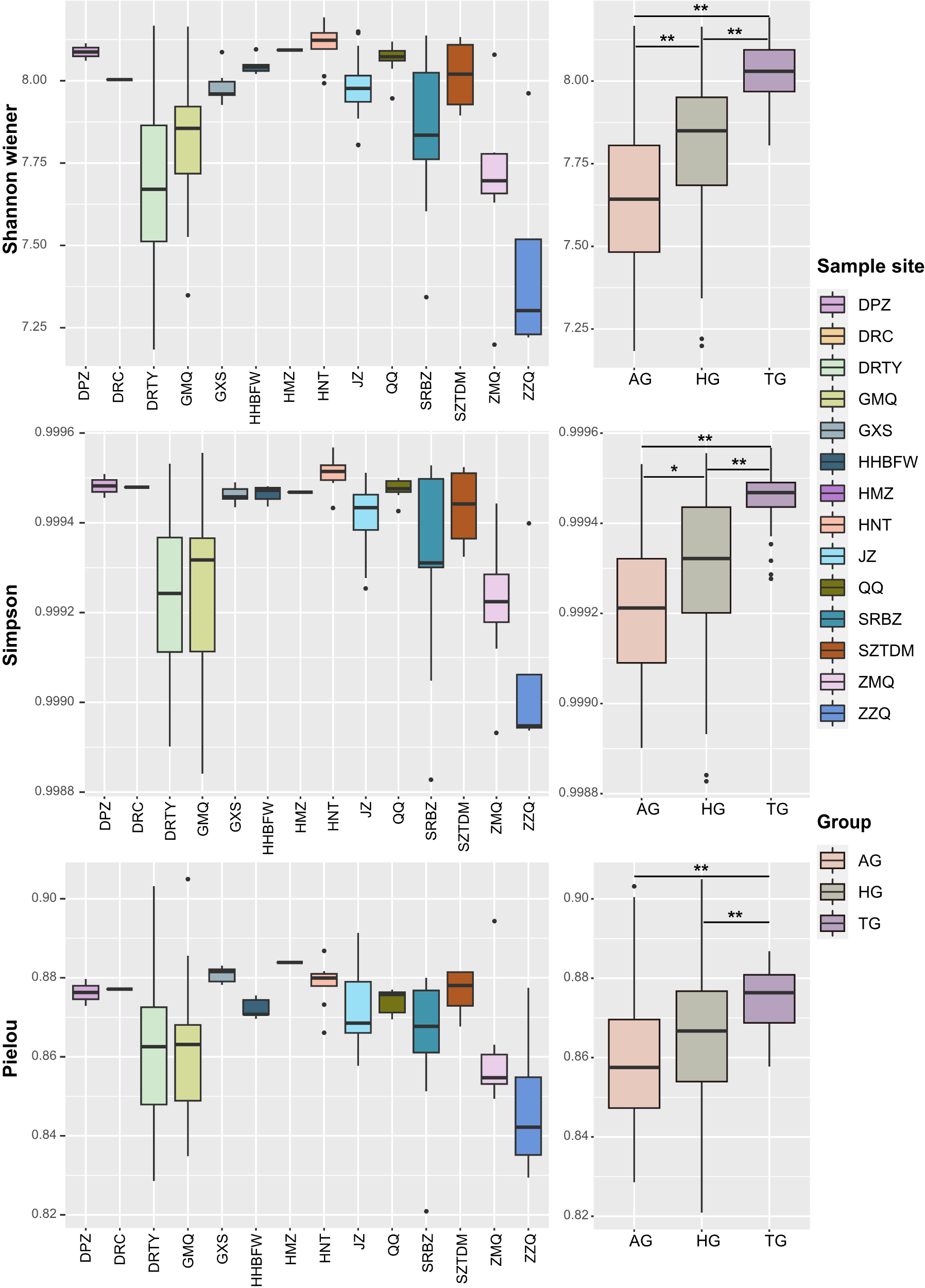

**Fig. S8. Boxplot of functional diversity of different sites and groups.** The significant difference among the three groups was tested using the Mann-Whitney *U* test. The “**” represented the significance ≤ 0.001.

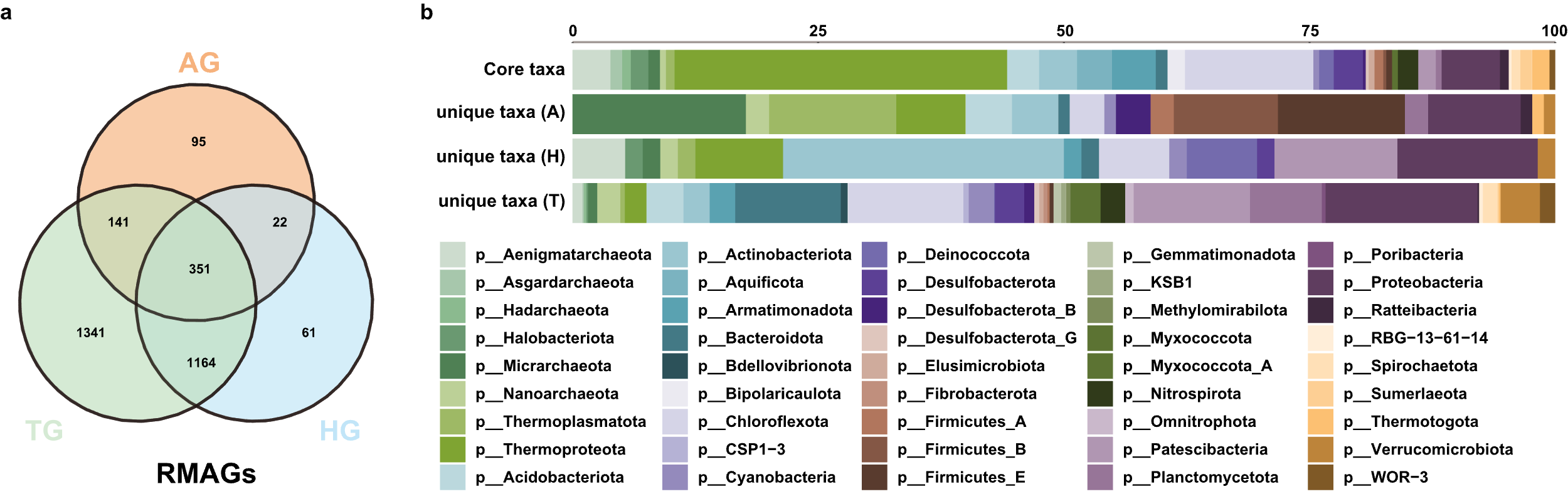

**Fig. S9. Unique and core species of the three groups. (a)** Venn diagram of species of three geothermal spring groups. **(b)** Unique and core species of the three groups. Different colors represented the different phyla.

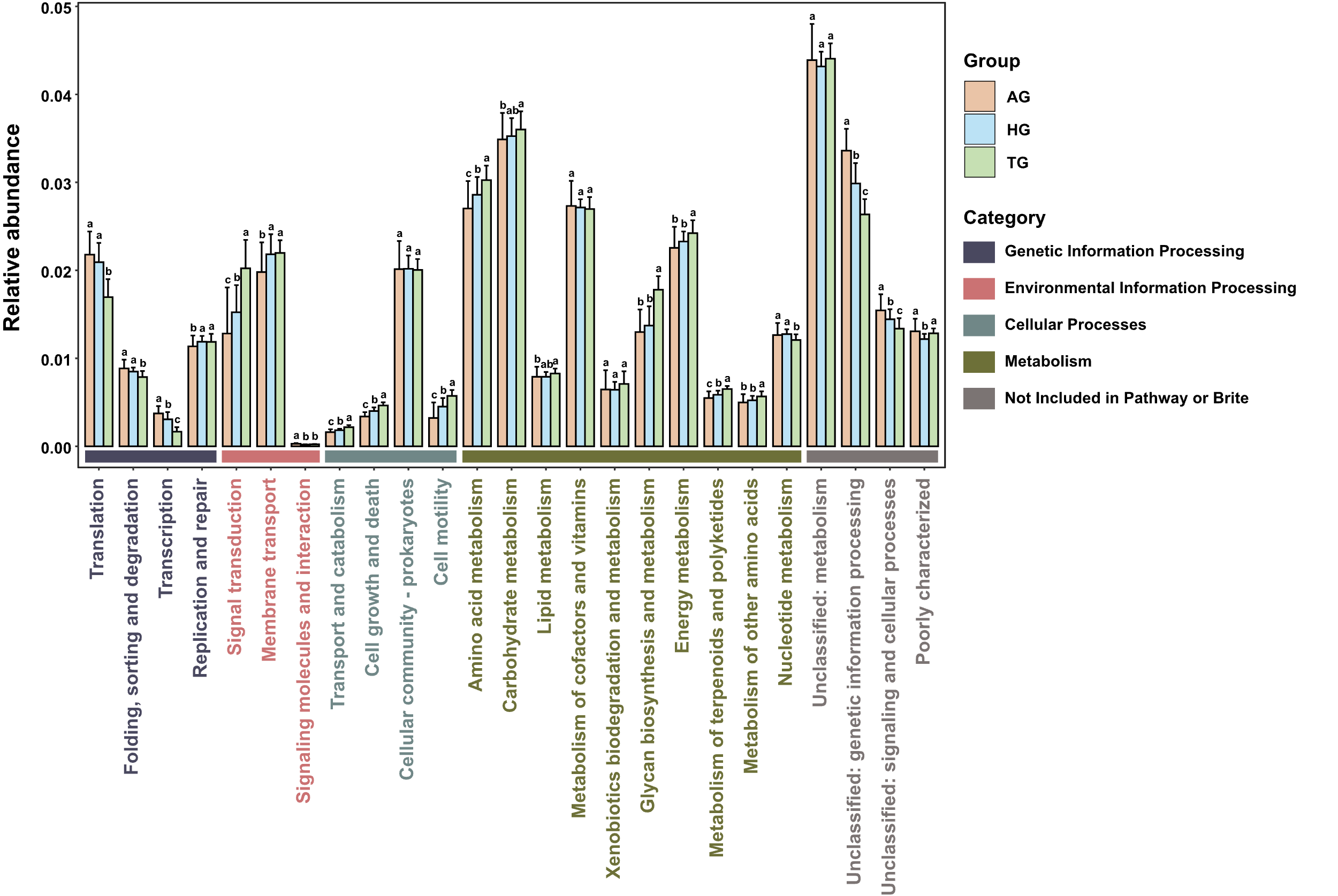

**Fig. S10. Metabolic category of accessory gene families of three habitats.** Based on annotation results of the KEGG database. The relative abundance of each metabolic category of the three groups was represented by bars with different colors. Differences among three habitat types in all categories were assessed using analysis of the variance (ANOVA, function “aov” in R package “agricolae”), followed by LSD post hoc all-pairwise comparisons test (function “LSD.test” in R package “agricolae”). Significant differences among groups were marked with alphabetic letters on the top of the bars.

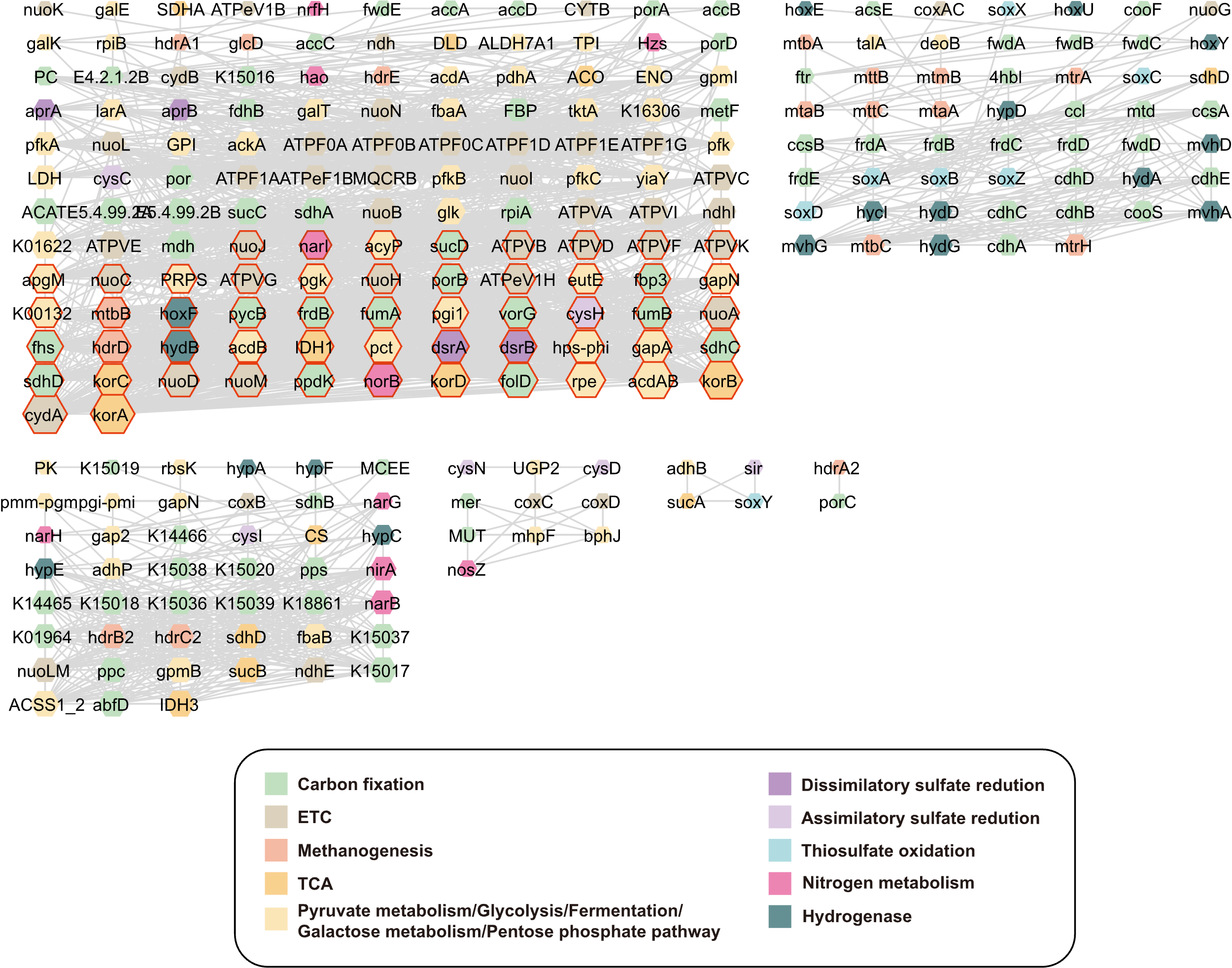

**Fig. S11. Co-occurrence functional networks of AG.** Functional genes of RMAGs within microbial networks of AG, involved in core metabolic pathways, were selected to construct the functional networks (details of these functional genes are provided in Table S16). The node size represented the nodes' degrees. Nodes with the top 50 degrees were marked with red strokes. Different colors indicated different metabolisms.

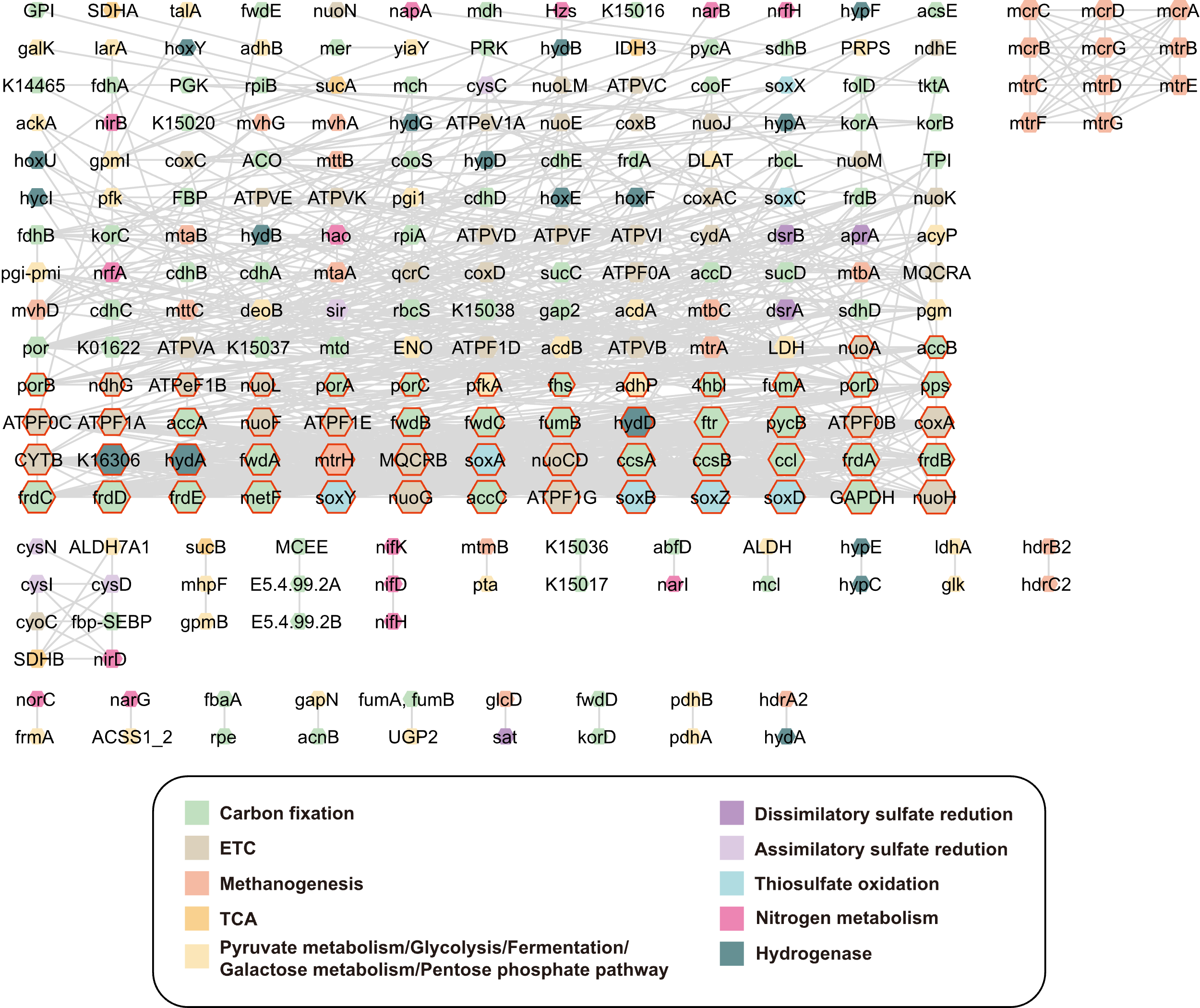

**Fig. S12. Co-occurrence functional networks of HG.** The corresponding information aligns with the networks of AG mentioned above.

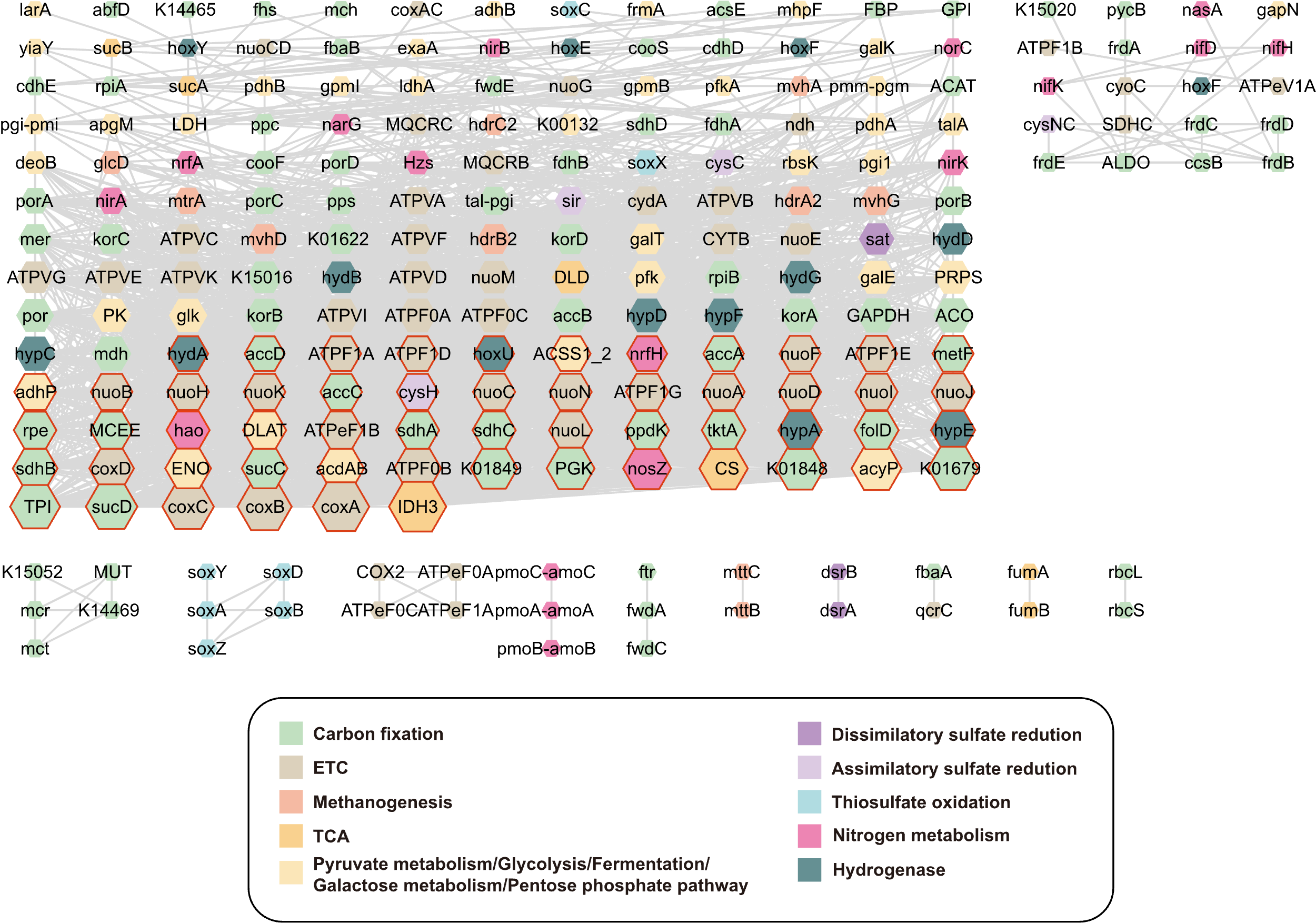

**Fig. S13. Co-occurrence functional networks of TG.** The corresponding information aligns with the networks of AG mentioned above.

**Table S1.** Geochemical characteristics of the 152 metagenome samples.

**Table S2.** Genome quality and characteristics statistics of all 12789 MAGs in this study.

**Table S3.** Mapping results of all 3176 RMAGs and assembly scaffolds over 500bp in 152 metagenomes based on BBMap.

**Table S4.** The relative abundance of all 3176 RMAGs in 152 metagenome samples.

**Table S5.** The relative abundance of archaea in 152 metagenome samples.

**Table S6.** The relative abundance of KOs in 152 metagenome samples.

**Table S7.** The relative abundance of KO categories in 152 metagenome samples.

**Table S8.** Mantel test between environmental variables and community composition.

**Table S9.** The alpha and functional diversity of 152 samples.

**Table S10.** The results of gene families in three hot spring groups.

**Table S11.** The relative abundance of accessory gene categories in 152 samples.

**Table S12.** The species network properties of three hot spring groups.

**Table S13.** Node information of RMAGs in three hot spring MENs.

**Table S14.** The functional network properties of three hot spring groups.

**Table S15.** Node information of KOs in three hot spring functional MENs.

**Table S16.** Metabolic potentials of RMAGs from three hot spring MENs.
